# Structural connectome dimension shapes brain dynamics in health and disease

**DOI:** 10.1101/2025.06.30.662336

**Authors:** Giacomo Barzon, Michele Allegra, Mohammad Hadi Aarabi, Lorenzo Pini, Manlio De Domenico, Maurizio Corbetta, Samir Suweis

## Abstract

The structural connectome serves as the foundation for neural information signaling, playing a primary constraint on brain functionality. Yet, its influence on emergent dynamical properties is not fully understood. Generally, a key measure of a system’s structural impact on dynamical phenomena is its dimension. By tracking the temporal evolution of diffusive perturbations, we estimate a scale-dependent measure of dimension of empirical connectomes. At the local scale, it is highly heterogeneous and follows a gradient from sensory-motor to high-cognitive areas. At the global scale, it encapsulates mesoscale topological information related to the balance between segregation and integration. Furthermore, by comparing connectomes from stroke patients, we find that dimension captures the local effects of lesions and, at a global level, is linked to impaired critical patterns and decreased cognitive performance. Overall, the dimension of the connectome may serve as a powerful descriptor for bridging the gap between structure and function in the human brain.

## I. INTRODUCTION

The intricate backbone of white matter connections – axonal tracts linking together distant gray matter regions – constitutes the so-called “structural connectome” of the brain [1]. It acts as the brain’s communication network, enabling signaling between neural populations and hence most of cognition, from basic sensory processing to complex cognitive tasks [2, 3]. Like many empirical networks, the structural connectome is highly heterogeneous, with broad distributions of connection strengths [4], hierarchical modular organization [5, 6], spatially dependent connectivity following the exponential distance rule (resulting in strong local and sparse long-range connections) [7], and diverse local connectivity motifs [8], with all these various topological properties intricately interconnected [9]. This non-random topological structure is thought to arise from an optimal compromise between signal processing demands and biological constraints (such as wiring cost) [10, 11] and to determine an optimal balance between segregation and integration [12–15].

The transfer of electrical and chemical signals in the brain is strongly shaped by the connectome’s topological properties, and previous research has highlighted the functional influence of global features (such as modularity or small-world propensity) and local ones (such as degree) [16]. However, dynamical processes evolving on complex networks are not determined by single, isolated topological properties, but are instead constrained by the geometry induced by the full connectivity structure [3, 17–19].

As for any dynamical process, emergent properties can be be strongly shaped by the dimensionality of the underlying connectivity space. In classical Euclidean spaces, dimensionality can be framed in simple geometric terms: it describes how the number of reachable points (found within a given distance from a reference point) increases as the distance grows. The rate at which this accessible set expands defines the dimension of the space [20]. Higher dimensionality implies that activity or signals can propagate along a larger number of distinct paths. Conversely, lower-dimensional spaces constrain propagation to fewer available directions. Consequently, dimensionality fundamentally shapes the dynamics of diffusion processes [21, 22] and the nature of phase transitions that can occur [23, 24], thereby favoring or hindering the emergence of criticality [25–28].

In real-world networks, which are characterized by highly heterogeneous structures, the notion of geometric distance is not uniquely defined. As a result, scaling relations based on spatial distance are not immediately generalizable, making the definition of dimensionality less straightforward [5, 29–38]. However, since the way diffusion unfolds depends systematically on the underlying dimension, one can infer dimensionality by measuring the network’s dynamical response to diffusive perturbation. This idea was recently operationalized in [39] by simulating the spreading of diffusive perturbations across the network. This method involves perturbing one node at a time and then observing how other nodes transiently respond to the perturbation over a defined period of time (Fig. 1a). This allows defining a local dimension of the network for the selected node [40] (Fig. 1b). The local dimension is a “mesoscale” metric, as it does not depend just on the nearest neighbors (“microscale”), nor on the structure of the entire network (“macroscale”). The variation of the local dimension across nodes provides insight into the network’s structural complexity and the way information propagates through its intricate architecture [41, 42]. Furthermore, a global measure of dimensionality can be obtained by averaging the local dimension over all nodes (Fig. 1c).

**FIG. 1.**
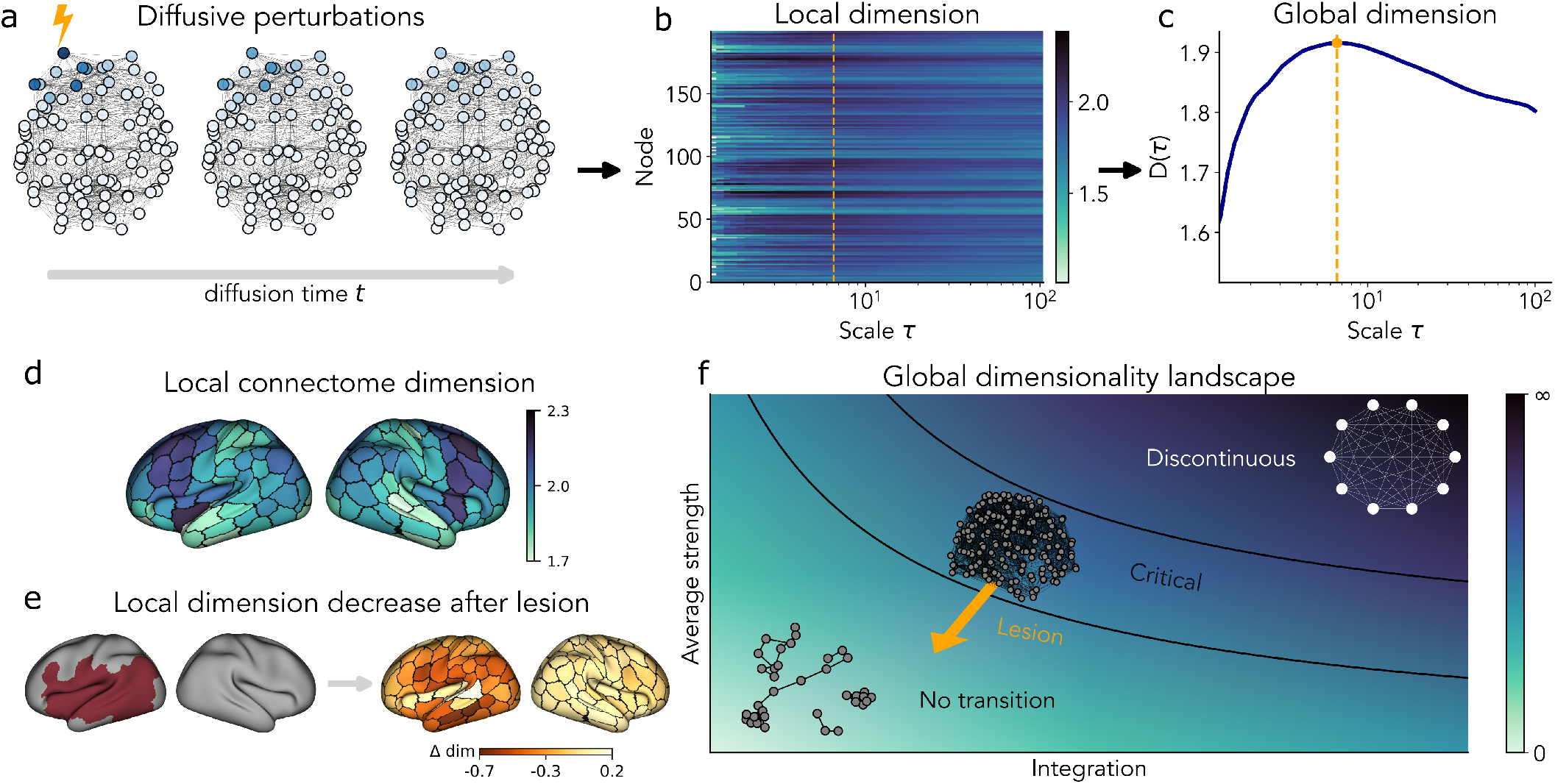
Multiscale dimensionality of the human connectome. (a) We track the spreading of diffusive perturbations across the connectome over time. (b) This temporal evolution allows us to estimate a local (nodal) measure of dimensionality at each scale *τ* (see Methods). (c) From the local dimensions, we derive a global measure of the network’s dimensionality. (d) The local dimension of the connectome is highly heterogeneous and aligns with a gradient from sensory-motor to associative areas. (e) The local dimension is influenced by focal lesions, leading to a decrease in areas near the lesion center. (f) The global dimension is influenced not only by overall connectivity (average strength), but also by how links are organized, reflecting the network’s integration. An optimal connectivity region is identified where critical features of the brain dynamics emerge. Focal lesions lead to reduced connectivity, impaired integration, and a loss of critical emergent features, overall shifting the connectome away from this optimal region which is captured by a decrease in the global network dimensionality.

In this study, we show that the dimension of the human connectome, both on the local and global scales, encapsulates key properties of the brain’s macroscopic organization (Fig. 1d). In particular, the local dimension of the connectome correlates significantly with the anatomical and functional organization of the brain, with higher dimensions found in the dorsal and medial regions involved in integrative functions. Additionally, local dimension is negatively correlated with myelin content, suggesting that areas involved in complex cognitive processes rely more on integrative connectivity than rapid, highly myelinated pathways. Instead, the global dimension is able to jointly capture seemingly disparate topological properties, including the network’s overall strength but also by mesoscale features such as community structure and topological heterogeneity, providing a comprehensive “holistic” topological indicator, directly related to the efficient transport of information in the brain. By focusing on the key example of stroke, we show that this indicator can be particularly suitable as a biomarker of structural damage in neurological disorders, which significantly affect the connectome and its properties [16, 43]. The dimension discriminates between healthy and damaged connectomes, quantifies the differential effect of the lesions in perturbing local networks centered around each brain area, and successfully captures the global impact of the lesion on information transport in the brain and cognitive impairment. Finally, by using a connectome-informed model of brain activity, we observe that the dimension is strongly related to the ability of the connectome to support critical dynamics, with drops in dimensionality accompanying the loss of critical dynamics observed in stroke patients [44].

## II. RESULTS

### A. Unraveling the effects of mesoscopic topological features on network dimensionality

We hypothesized that the non-random structure of complex networks directly influences their dimensionality. To this aim, we examined both the impact of the overall connectivity level (average strength), and non-random topological features that are commonly observed in brain connectomes. These features encompass the balance between integration and segregation, the degree of order versus disorder, and the heterogeneity of link strength within the network (Fig. 2). Specifically, we generated synthetic networks with different levels of topological disorder using the Watts–Strogatz model [45] by varying the probability of rewiring, with different levels of community structure, using the stochastic block model [46] and varying the ratio between inter- and intra-community connections (see Methods). Recognizing that non-random structure involves not only the distribution of connections but also their strengths, we incorporated this aspect in our analysis by assigning weights to the network links. Since connectivity strengths in the cortex are well established to follow a Lognormal distribution [4, 47, 48], we generated connection weights from a Lognormal distribution by changing its variance (see Methods).

**FIG. 2.**
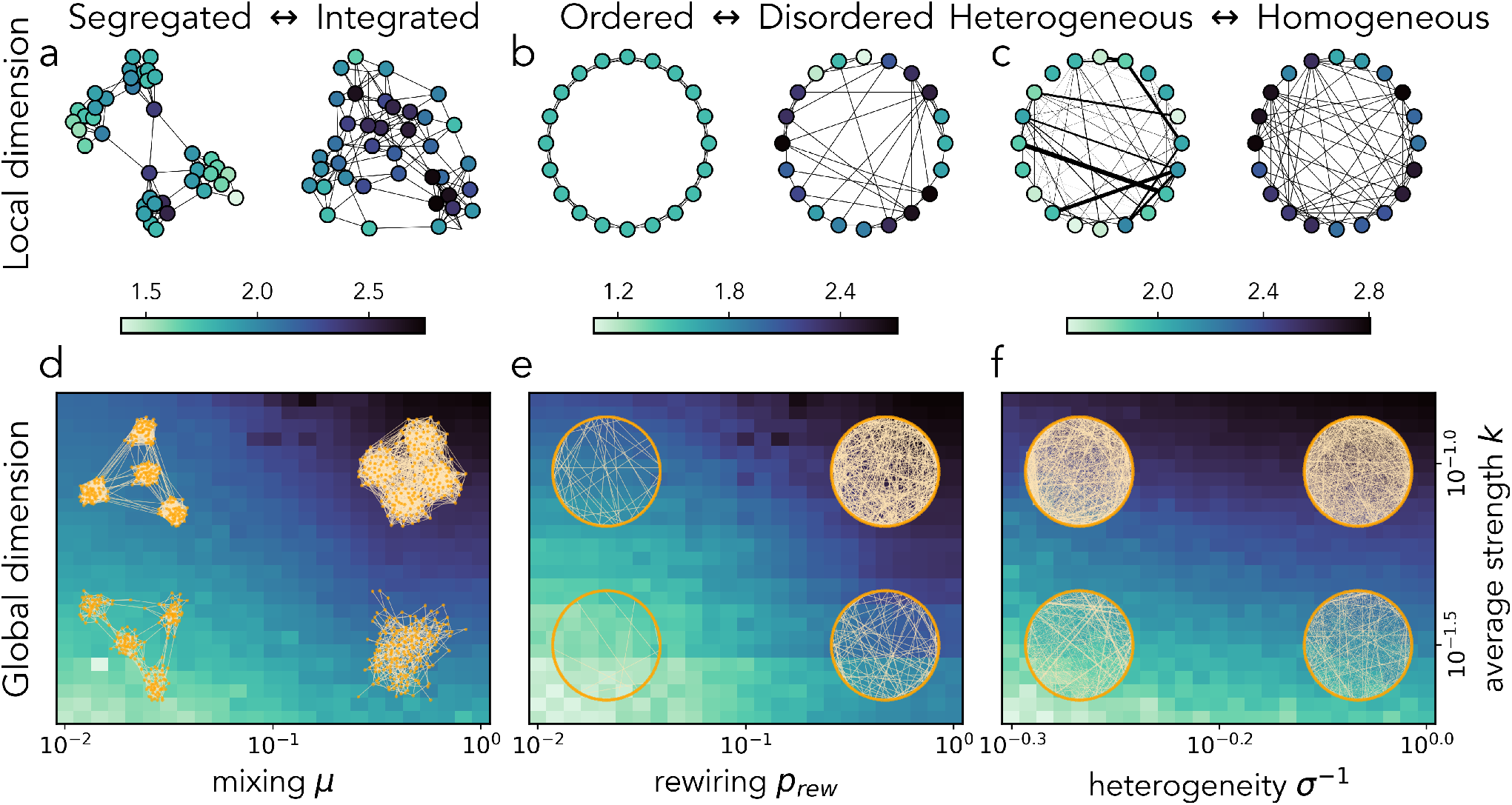
Effects of topological features on network dimensionality. (a) Segregation versus integration, modeled by stochastic block networks with four communities, for different mixing parameters *µ*. (b) Order versus disorder, modeled by Watts–Strogatz small-world networks with different rewiring probability *p*_*rew*_. (c) Weight heterogeneity versus homogeneity, modeled by an Erdős-Renyi random network with weight generated from a lognormal distribution with same mean but different variance *σ*. To ensure a consistent visualization, we plot the inverse of *σ*, since dimensionality increases as heterogeneity decreases. In (a-c) we compute the local dimension, while in (d-f) we compute the global dimension by varying also the average strength *k*. The colorbar for each column represents the values of both the local dimension (top row) and the global dimension (bottom row) for the corresponding network ensemble.

By systematically adjusting these parameters, along with the overall average strength, we were able to observe how changes in network topology affect dimensional metrics. At the local level, we found that dimension is often highly heterogeneous (Fig. 2a-c). For example, nodes positioned between two communities tended to have a larger dimension than nodes connected within a single community (Fig. 2a). This suggests that nodes from which one can easily reach different parts of the network have a higher local dimension [39, 49]. Similarly, more connected nodes displayed a larger local dimension (Fig. 2b-c). Our analysis thus suggested that the dimension is larger for more central nodes, which is consistent with previous studies [39, 49].

While local dimensions provide valuable insights into the role of individual nodes within their network surroundings, they do not fully capture the network’s overall structural organization or its capacity for global information integration and communication. Therefore, we extended our analysis to the global scale by computing the average of local dimensions across all nodes. This global dimension serves as an integrated measure that encapsulates the network’s overall potential in supporting efficient information transfer. We found that average strength positively correlates with the global dimension in all classes of networks (Fig. 2d-f). We also found a non-random topological structure that significantly influences the global dimension of a network: segregation or integration. Highly segregated networks exhibited low dimensions, which increase with a higher probability of intracommunity links (Fig. 2d). A similar effect was observed when increasing the number of shortcuts in a regular ring structure (Fig. 2e). Conversely, heterogeneity in connection weights tended to decrease the global dimension of a network (Fig. 2f).

Overall, our findings indicated that network dimensionality effectively captures the influence of mesoscopic topological features on the network organization, both on local and global scales.

### B. Variation of the local dimension across the human brain connectome

After establishing that the local dimension effectively captured information about the hierarchical and central position of nodes within synthetic networks, we turned our attention to healthy human brain connectomes to examine whether the local dimension of brain regions within the whole-brain network is associated with the regions’ microstructural features and their functional role.

We thus considered structural connectivity data from 33 healthy participants, parcellated according to the Schaefer brain atlas [52] (200 cortical regions, see Methods). We computed the local dimension for each individual connectome and we averaged the results across all subjects for each cortical region (Fig. 3a). We found that the local dimension exhibits a significant correlation with the brain’s anatomical organization. Specifically, we observed that dorsal regions tend to have a higher local dimension, which progressively decreases as we move toward ventral ones (Fig. 3b, *ρ* = 0.55, *p <* 0.001). Similarly, we found a gradient along the medial-lateral axis, with the local dimension being higher in medial regions and decreasing laterally (Fig. 3c, *ρ* = −0.35, *p <* 0.001). Instead, we did not find any relationship with the anterior-posterior axis, nor did we observe any left-right asymmetry (see Supplementary Information, Fig. S1).

**FIG. 3.**
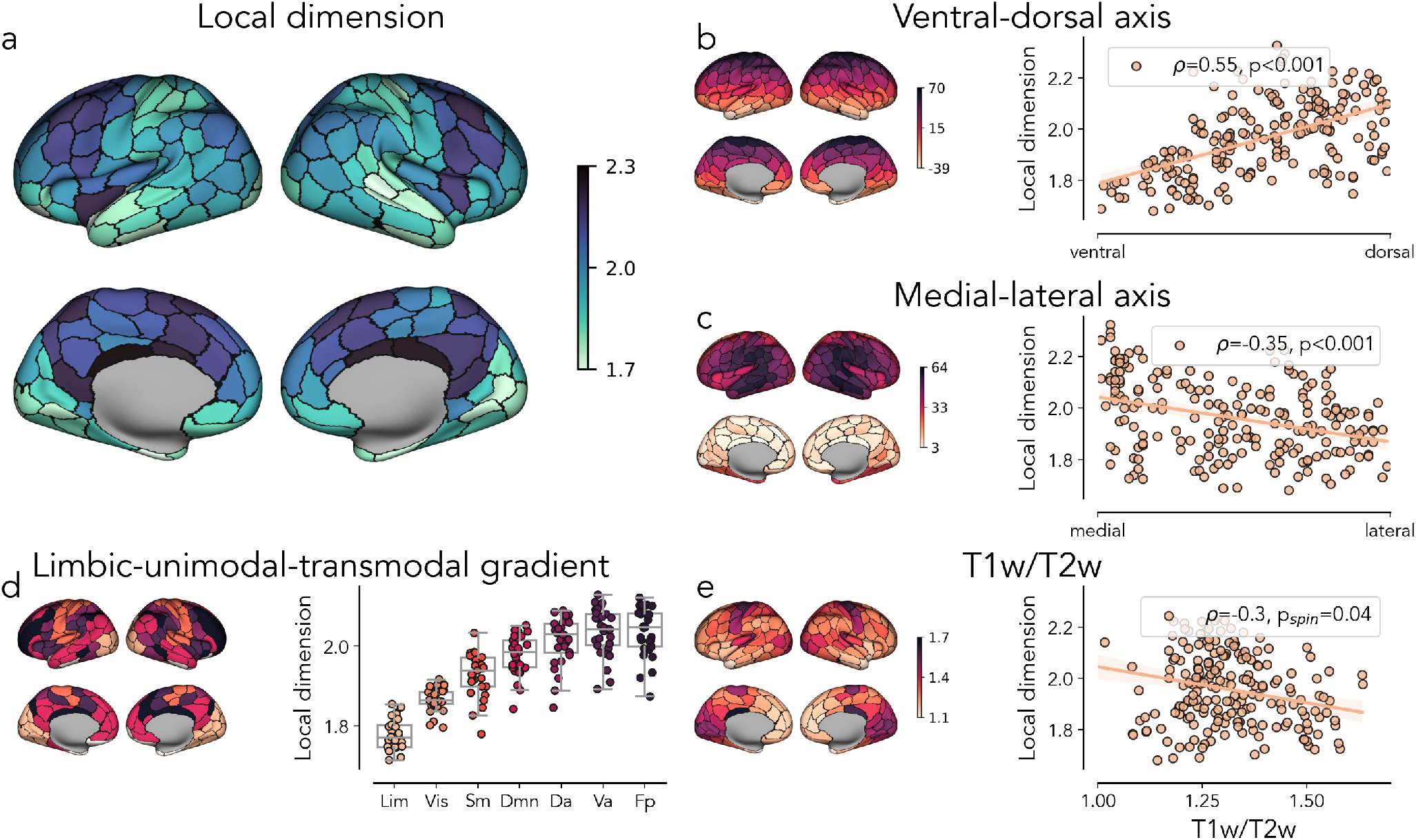
Organization of local dimension in the human brain connectome. (a) Average local dimension across all control subjects. (b) Relation between local dimension and ventral-dorsal axis. (c) Relation between local dimension and medial-lateral axis. (d) Average local dimension within canonical resting-state networks. (e) Relation between local dimension and regional T1w/T2w ratio [50], that is publicly accessible via the *neuromaps* toolbox [51]. The single connectomes and associated maps were parcellated using the Schaefer 200 parcellation scheme [52].

We then examined the relation between the regions’ dimension and their membership in the canonical resting-state networks, which are well-established functional communities [52]. By averaging the local dimensions of regions belonging to each functional network, we found that transmodal regions, such as those involved in the default mode and frontoparietal networks, exhibited significantly higher dimensions compared to unimodal regions, like those in the visual and somatosensory networks (Fig. 3d). Overall, the ventral-dorsal, the lateral-medial, and the limbic-unimodal-transmodal gradients of the local dimension, may reflect the fact that regions more involved in higher-order processing (which belong to transmodal networks and are prevalently dorsal and medial) have more complex connectivity patterns allowing them to effectively engage in long-distance communication with many other areas to sustain their integrative role.

This hypothesis was further supported by examining the relationship between local dimension and the T1w/T2w ratio, a measure commonly used as a proxy for cortical myelination [50]. Previous studies have demonstrated that myelin distribution in the brain is not uniform, as regions associated with higher cognitive functions typically have lower levels of myelin compared to primary sensory and motor areas [50, 53]. It is widely held that this variation reflects a lower reliance on areas involved in complex, integrative processes with highly myelinated connections allowing for rapid communication. Consistent with the literature, our analysis showed that regions with higher local dimensions tend to have smaller T1w/T2w ratio (Fig.3e, *ρ* = −0.30). The negative correlation remained significant after performing a spin test (*p*_*spin*_ = 0.04, *N*_*spin*_ = 10^4^; see Methods), indicating that the relation is not trivially explained by spatial autocorrelation in the two maps.

Importantly, these spatial and functional patterns are significantly disrupted when the local dimensionality is computed on strength-preserving, shuffled versions of the connectome obtained through Maslov-Sneppen weight-shuffling or degree-preserving randomization [54] (Supplementary Information, Fig. S2). These finding corroborates the specificity of the relationship between local dimensionality and the explored brain features, demonstrating that the observed dimensionality maps are not merely a reflection of nodal degree or strength but are rooted in the non-random, higher-order topological organization of the human brain.

### C. Connectome dimensionality captures structural and cognitive alterations after focal lesions

While the local dimension offers insights into regional variations within the brain, the global dimension can shed light on the overall structural organization of individual connectomes, potentially highlighting the impact of pathological conditions. To investigate this, we compared the global dimensions of connectomes from stroke patients with those from healthy controls (see Methods). We observe a significant reduction in global dimension for stroke patients compared to healthy controls (Fig. 4a, independent samples t-test: *t* = −4.01, *p <* 0.001). Unsurprisingly, this decrease became more pronounced for subjects with larger lesions (Fig. 4b, *ρ* = −0.31, *p* = 0.03), supporting the intuition that more extensive brain damage leads to a greater impact on the overall structure of the network.

**FIG. 4.**
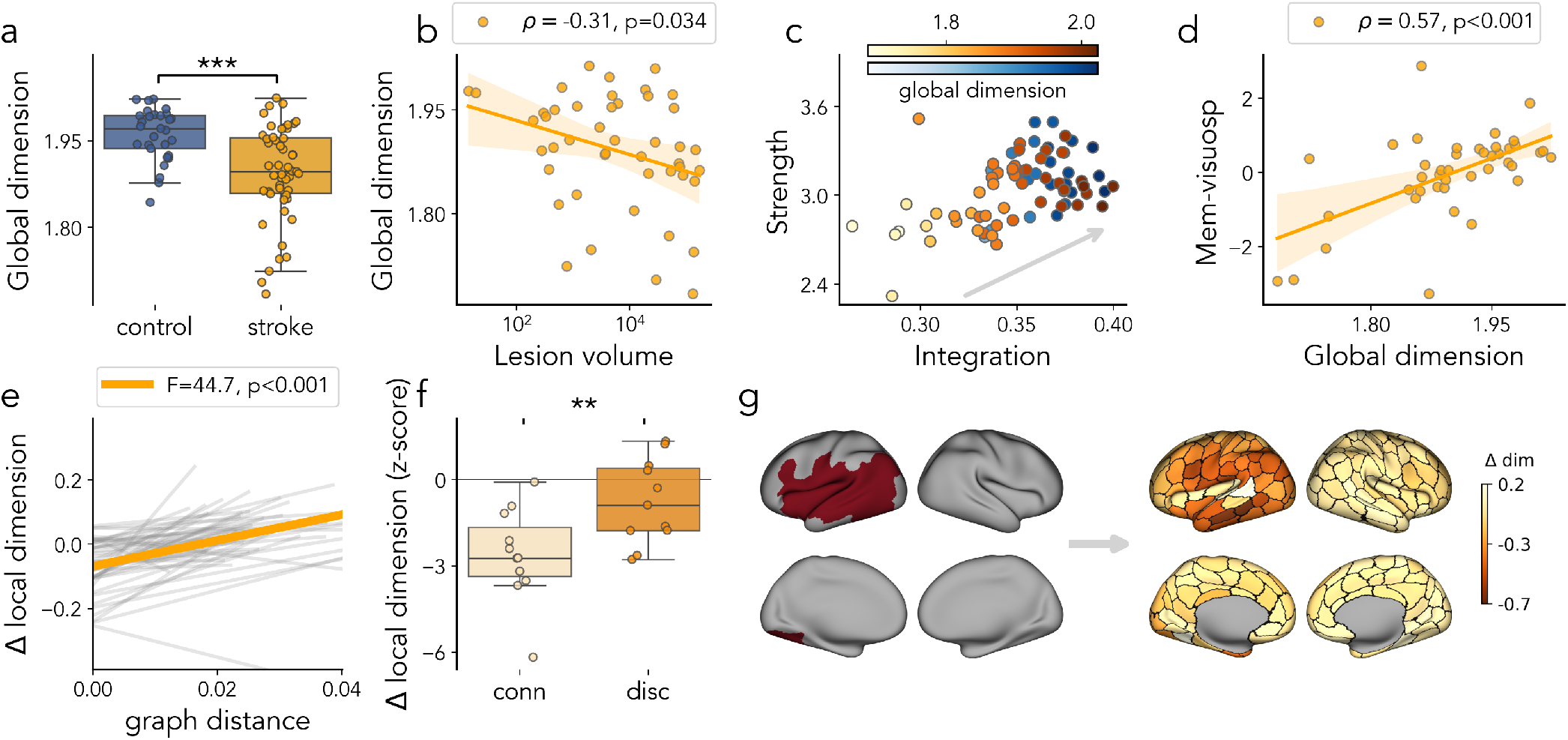
Local and global dimension in pathological connectomes. (a) Decrease in network dimension in stroke patients compared to controls. (b) Relation between dimensionality in stroke connectomes and lesion volume. (c) Relation between dimensionality and structural features. Points are color-coded according to their global dimension, with warmer colors indicating higher dimensionality. An arrow is overlaid to highlight the principal trend in the data, showing that connectomes with both higher integration and greater average strength tend to exhibit higher global dimensionality. (d) Relation between dimensionality and cognitive abilities in stroke patients. (e) Relation between local reduction in dimensionality and distance from the lesion (computed as the shortest network path to the lesion center), as shown by the random, subject-wise (gray), and fixed, group (orange), effects. (f) Decrease in local dimension for connected and disconnected regions. (g) Example of lesion mask and local reduction in dimensionality for one patient.

In patients, we observed a positive correlation between the global dimension and the total network connectivity, as measured by the average strength (Fig. 4c, *ρ* = 0.54, *p <* 0.001). A weaker relation was observed in healthy controls (*ρ* = 0.34, *p* = 0.07). This suggests that the dimension decrease observed in patients can be partially explained in terms of simple disconnections.

Nevertheless, the reduction in global dimension cannot be solely attributed to a decline in overall connectivity. In particular, the brain’s capacity for integrated communication across regions plays a critical role in shaping large-scale dynamics. We captured this property using a measure of network integration, defined as *I* = 1 − *Q*, where *Q* is the modularity of the structural connectivity graph (see Methods). Higher integration reflects a network that is less segregated into isolated communities and better supports global communication. We found that individual differences in global dimensionality were more strongly associated with variations in integration than with total connection strength (Fig. 4c), both in healthy controls (*ρ* = 0.89, *p <* 0.001) and in stroke patients (*ρ* = 0.96, *p <* 0.001). This result was robust with respect to the community detection method used (Supplementary Information, Fig. S3). This relationship remained significant even after accounting for the effect of average strength through partial correlation analysis (control: *ρ* = 0.95, *p <* 0.001; stroke: *ρ* = 0.96, *p <* 0.001; see Methods). This suggests that the observed reductions in dimensionality are shaped not only by the extent of disconnection, but also by how those disconnections alter the brain’s integrative architecture.

As the observed global dimensionality reduction was associated with focal lesions, we hypothesized that it corresponds to a pattern of local reductions, with regions closer to the lesion exhibiting more pronounced decreases. To test this hypothesis, we calculated the change in dimension for each region in stroke patients relative to the average local dimension in healthy controls. We then used a linear mixed model, incorporating subjects as a random factor for both slope and intercept, to assess the impact of a region’s graph distance from the lesion center (computed as the weighted shortest path distance) on its dimensionality (Fig. 4e, see Methods). We found that it significantly affected the local reduction in dimensionality (F test, *F* = 44.7, *p <* 0.001), with a stronger effect than the spatial (Euclidean) distance (F test, *F* = 6.76, *p* = 0.012; see Supplemental Information, Fig. S4). Thus, reductions in dimensionality were more pronounced in regions closer to the lesion, and this effect decreased with increasing distance. Notably, the network (graph) distance from the lesion provided a better explanation of this pattern than simple spatial (Euclidean) distance, underscoring that the impact of a lesion propagates primarily along the brain’s structural connectivity rather than through physical proximity alone at the level of individual subjects (Fig. 4g).

To further investigate the role of structural disconnection in shaping local reductions in dimensionality, we compared the average dimensionality change between regions that remained connected and those that became disconnected following stroke (Fig. 4f). Disconnected regions were defined as those whose connection strength to the rest of the network was more than two standard deviations below the mean network strength observed in healthy controls. This analysis revealed that the decrease in local dimensionality was significantly more pronounced in disconnected regions (independent samples t-test: *t* = − 12.2, *p* = 0.006), supporting the hypothesis that the pattern of structural disconnection critically contributes to the observed reductions in dimensionality.

Given that the global dimension reflects broad alterations in network organization and the extent of focal damage caused by stroke, we hypothesized a direct relation between the global dimension and cognitive impairment. To test this, we examined its association with behavioral factors previously identified via factor analysis of a comprehensive neuropsychological battery administered to the same cohort [55] (see Methods). This analysis identifies underlying latent variables that summarize key cognitive domains, such as memory, visuospatial abilities, and executive function. We found that a lower global dimension was associated with more pronounced cognitive deficits, as identified by a factor related to memory and visuospatial abilities (Fig. 4d). This relationship was observed exclusively in stroke patients, as no significant correlation was found in healthy controls, nor in the other behavioral factors (see Supplemental Information, Fig. S5). This finding underscores the functional significance of network dimensionality in supporting cognitive abilities and highlights the potential of global dimension as a biomarker for stroke-related cognitive impairments.

To test the robustness of all these findings, we also computed the spectral dimension of the connectomes (see Methods and Supplemental Information, Fig. S6-S8), confirming that the observed decrease in connectome dimension is consistent across different estimation methods of network dimensionality.

### D. Connectome dimensionality affects criticality

Finally, we investigated how the global dimension of the structural connectome affects large-scale dynamics. To do so, we implemented a personalized whole-brain model using the corresponding individual connectome as input. Specifically, we simulated a variant of the Greenberg-Hastings cellular automata with homeostatic normalization [44, 56–58] (Fig. 5a, see Methods). The dynamics of this model is notably influenced by the threshold parameter *T*, which regulates the system’s excitability and serves as a key control parameter.

**FIG. 5.**
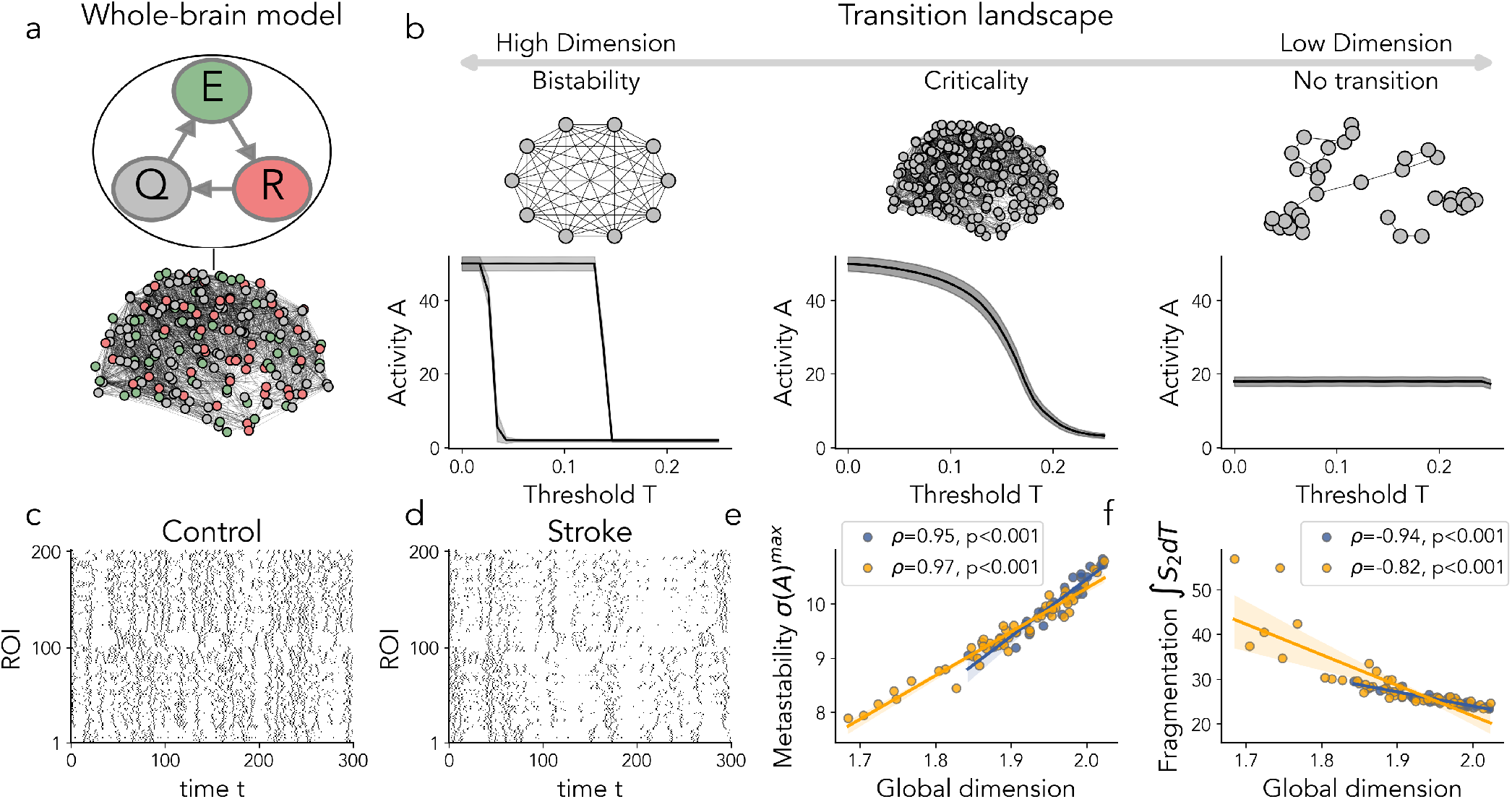
Connectome dimension and its relation with emergent dynamical patterns. (a) Stochastic whole-brain model [44, 56–58]. Shaded regions represent the standard error across independent simulation runs (see Methods). (b) Network dimension predicts transition type. (c,d) Example of simulated trajectories from a healthy and stroke connectome (T=0.16). (e) Relation between connectome dimensionality and metastability. (f) Relation between connectome dimensionality and fragmentation.

We compared the average activity levels — defined as the average number of active nodes (see Methods) — across different threshold levels for network topologies with different global dimensions. Specifically, we analyzed a typical healthy connectome alongside two extreme cases: a fully connected network, which is characterized by an infinite global dimension in the thermodynamic limit, and a disconnected network, which corresponds to a null global dimension (Fig. 5b).

In a fully connected network, the dynamics exhibited a bistable phase, aligning with the analytical predictions from mean-field analysis [58]. In a completely disconnected network, the activity remained low for all values of *T*. In contrast, when using the human connectome, the dynamics exhibited symmetry-breaking phenomena leading to a continuous (critical) transition [56]. This suggests that connectomes correspond to an “optimum dimension” — shaped by various mesoscale features — that is instrumental in facilitating the emergence of critical dynamical properties. If the dimension is reduced, as in stroke patients, this may lead to a loss of critical properties. To show this, we simulated the model using both healthy and stroke connectomes. The trajectories of neural activity in healthy brains exhibited a high degree of complexity and variability (Fig. 5c). In contrast, the trajectories for stroke patients showed more stereotyped patterns with distinct periods of sparse activation (Fig. 5d). From the simulated trajectories, we computed several key observables that typically peak at intermediate values of *T* corresponding to the critical transition (see Supplementary Information, Fig. S9). For instance, we calculated the average activity variance at the critical point, often referred to as “metastability” [59, 60], which reflects the degree of variability and fluctuations in neural activity. Additionally, as done in [44], we assessed the integral of the second cluster size over all the threshold values, that we termed “fragmentation” (see Methods). While the largest cluster changes monotonically, transitioning from an integrated active core to a completely deactivated one, the second largest cluster captures the formation of major separate active components. By integrating this value, we quantify the total extent to which the system’s activity is distributed among these disconnected groups. Our analysis revealed a significant correlation between the values of these metrics and the dimensionality of the connectome. Specifically, a reduction in the dimensionality of stroke connectomes was associated with diminished metastability (Fig. 5e) and increased fragmentation (Fig. 5f), indicating that anomalous activity patterns unfolding over connectomes with a reduced network dimensionality were associated with a loss of critical properties.

## III. DISCUSSION

The large-scale organization of the brain reflects the tension between local processing and long-distance communication, i.e., segregation and integration. At a structural level, this tension results in a structural connectome characterized by highly non-random features (clustering, modularity, and the small-world property) and a hierarchical structure where integration is predominantly exerted by associative regions. At a functional level, the tension reflects in dynamics poised between order and disorder, presenting many of the characteristics of so-called critical phenomena. In this work, we showed that a single metric, the network’s dimension, can capture both the *local* balance between segregation and integration in different brain areas, and the *global* balance struck by the connectome as a whole. Furthermore, this metric sheds light onto the structure-function relation in the brain: only a small range of dimensions will allow the connectome to support critical-like dynamics. Alterations of the structural connectome caused by brain pathologies typically disrupt the healthy segregation/integration balance, both at the local and global level, leading to dramatic dynamical anomalies. By focusing on a prototypical case of brain pathology, stroke, we showed that dimension alone can suffice to track the occurring shift in the integration/segregation balance and the consequent loss of critical features in the dynamics. Thus, dimension arises as a powerful marker of brain pathology. Notably, we replicated this analysis using both coarser (100 nodes) and finer (500 nodes) parcellation schemes, confirming the robustness of our findings (see Supplemental Information).

### A. Dimension of a complex network

In recent decades, simulating the response of diffusive dynamics has emerged as a powerful approach to estimate network properties that go beyond trivial, first-order geometric features [61, 62]. The time horizon of the diffusion acts as a natural scale factor: as activity spreads from a node, it progressively explores the network along multiple, heterogeneous pathways, capturing how the structure constrains and channels signal propagation [42, 49, 63]. This approach can be extended for estimating the dimension of a complex network [39]. An optimal scale naturally emerges as the peak of the global dimension: while in Euclidean lattices this is the scale that reproduces theoretical predictions, in empirical connectomes it captures a scale long enough to move beyond purely local features, yet short enough to maintain sensitivity to structural details that are averaged out at the global scale. This approach to measure dimensionality is conceptually similar to the spectral dimension [30, 35, 36, 63], in that it relies on the spreading of diffusive perturbations to define the dimension of a complex network. However, the framework introduced by [39] allows the dimensionality to be defined at the local (nodal) level. The concept of global dimension provides important characteristics about the embedding space and dynamical process on networks [5, 20, 22, 28, 30, 35, 36, 64], but substantially richer information can be obtained by characterizing the dimension locally [39, 40], enabling a multi-scale characterization of the network that global measures cannot provide. Importantly, comparative analyses revealed that local dimension captures unique topological information compared to classical centrality measures (see Supplemental Information, Fig. S11), and comparisons with randomized null models confirmed that the observed spatial gradients and global organization are not mere reflections of nodal degree or strength, but are rooted in the non-random architecture of the human connectome (see Supplemental Information, Fig. S2).

### B. Dimension and the integration/segregation balance

The organization of the brain connectome, not only in primates but across species, is shaped by two major forces: the need to minimize physical and metabolic costs, and the necessity to maintain efficient communication [16, 65, 66]. Cost minimization encourages the formation of dense, localized connections between nearby neural elements, resulting in compact network modules that facilitate specialized neural processing [67–69]. On the other side, the brain has evolved to form long-range connections, which, though costly, are essential for promoting global communication and functional integration across spatially and functionally distinct brain regions [70–72]. These long-range connections support a small-world topology, with a decrease in the average path length between nodes.

The competition between these forces determines an optimal dimension, both at the local and global levels. Locally, all nodes in the connectome exhibit a dimension between 1.7 and 2.3 (Fig. 3a), values that are broadly compatible with the notion that the cortex is organized as a folded surface with an intrinsic dimension close to 2, but with regional variations [73]. These local differences reflect a “segregation-integration” gradient, revealing a clear distinction between sensorimotor processing networks (SMN/VIS) and high-cognitive functions (DAN/VAN/DMN) (Fig. 3d). Sensorimotor regions, which are generally more segregated and modular, typically exhibit lower dimensions. Limbic regions, which are relatively isolated and specialized in emotional processing, memory, and basic survival functions, also show lower dimensions. Instead, associative regions, involved in executive functions and cross-modal integration, exhibit higher local dimensions. These high-dimensional regions are predominantly dorsal and medial, contributing to the observed organization of dimensional variation along a ventral-dorsal and medial-lateral axis (Fig. 3b-c). Notably, this axis does not align with the well-known principal gradient of functional connectivity [74], which is not primarily a segregation-integration axis, as it mainly distinguishes between regions devoted to external (SMN, VIS, and DAN) versus internal (DMN) cognition. The stronger/weaker integrative role of different regions also explains the observed anticorrelation between the local dimension and the myelin map (Fig. 3e): primary networks tend to be more myelinated because they handle immediate sensory input and motor control, requiring fast, reliable communication to ensure rapid responses to the external environment; in contrast, higher-order association networks are less myelinated, as they rely more on complex, integrative processing than on rapid signal transmission [53]. In future work, it will be interesting to compare the local dimension results in humans with other non-human primates connectomes: as associative regions have undergone significant expansion during cortical evolution [75], we expect to observe an increase of their local dimension in humans.

Globally, the connectome as a whole also strikes an optimal balance between segregation and integration. This can be visualized in the so-called “connectivity morphospace” [65, 66], an abstract space defined by a set of independent topological features, where each network occupies a specific location. In Figure 1f we see a simplified morphospace defined by two features, integration and average strength. “Optimal” connectomes occupy a limited region of this space, characterized by intermediate dimensions. Through experiments with synthetic networks, where we manipulated non-random network features (modularity, the small-world property, and weight heterogeneity), we showed that all these features influence the global dimension (Fig. 2d-f). We can thus hypothesize that most of the previously observed non-random features of the connectome collectively shape the observed network’s dimensionality, thus concurring to determine the optimal segregation/integration balance of the network. This hypothesis is supported by evidence of a strong correlation between dimensionality and the level of integration in individual connectomes (Fig. 4c).

### C. Dimension and critical dynamics

In recent years, the brain criticality hypothesis has gained traction as a framework to explain the emergence of global functional patterns in brain dynamics [76–80]. This hypothesis suggests that the brain operates near a critical point, balancing between order and chaos, which enables optimal information processing, flexibility, and responsiveness to stimuli [81–83]. The non-random topological features of the connectome – such as structural disorder, long-range connections, and hierarchical modularity – seem to play a crucial role in the emergence and maintenance of criticality [5, 6, 58, 84]. Since all these seemingly disparate features jointly concur in determining the integration/segregation balance, summarized by the network’s dimension, one may conjecture a strong relation between dimension and criticality.

To investigate such relation, we employed a well-established whole-brain model [44, 56–58] (Fig. 5a). Despite its simplicity, this model has proven effective in linking topology with functional patterns both in healthy individuals [56, 57] and in pathological conditions [44]. In networks with higher dimensionality, like fully-connected graphs, the model exhibits two stable equilibria, leading to a region of bistability with an associated discontinuous (noncritical) transition, in agreement with analytical predictions obtained in the mean-field limit [58]. In weakly connected networks, it yields a single stable phase characterized by low activity. Between these two extremes, the model can feature symmetry breaking and the emergence of critical transitions [56]. This phenomenon mirrors effects extensively studied in the context of the physics of impure materials, where the nature of phase transitions can shift from discontinuous to critical in the presence of quenched disorder. Crucially, this shift occurs only below a certain dimension [25–27].

The topology of the human brain connectome is poised toward this critical regime [56]. Our analysis suggests that this occurs because the connectome topology lies within a specific dimensional range that supports criticality and complex spatiotemporal activity patterns (Fig. 5b-d). Therefore, our findings strongly support the idea that the dimensionality of the underlying structural topology is a crucial factor in reconciling the bistable nature of activity with criticality through a symmetry-breaking mechanism [85, 86].

Comparisons with synthetic random networks performed in previous works [58, 87] have highlighted the specificity of the critical regime, demonstrating that critical transitions only arise within a narrow range of low degree and limited rewiring, closely matching the human connectome; outside this range, loss of criticality occurs, yielding either discontinuous or stable dynamics. Our observations allow us to reinterpret previous results, both in empirical and synthetic networks, in terms of the network’s dimensionality, providing a unifying framework to understand the conditions under which critical dynamics emerge.

To further assess the specificity of the observed dynamics beyond degree statistics alone, we performed additional simulations using degree-preserving randomized connectomes (Supplemental Information, Fig. S12). Degree-preserving rewiring significantly alters the dynamical order parameters while preserving signatures consistent with a critical transition, despite increasing global dimensionality (Supplementary Fig. S2e–f). These findings are consistent with previous results obtained in synthetic networks [58], where imposing empirical weight structure onto random connectivity restored the critical regime. Together with our results in Fig. 2c,f, this suggests that heterogeneity in the connectivity weights contributes to reducing network dimensionality relative to homogeneous networks, thus fostering the emergence of criticality.

Structural lesions in stroke patients act as a “natural” perturbation of network topology. The loss of critical dynamics in stroke patients, previously traced back to specific topological changes (average connectivity and integration) [44], can be clearly explained in terms of a reduced global dimension, demonstrating the relevance of dimensionality in shaping network dynamics.

Notably, the emergence of continuous transitions at intermediate dimensionality is not unique to the specific dynamical model we employed. Similar behavior has been reported in other neural dynamics models in which node activation depends on the integrated input from connected neighbors [5, 28, 64]. This convergence across models reinforces the idea that the link between dimensionality and criticality is a general feature of dynamical systems governed by integrative activation mechanisms, rather than a model-specific effect.

### D. Dimension and brain pathology

In stroke patients, we consistently observed a reduction in global network dimensionality as compared with healthy subjects (Fig. 4a). This reduction is not a mere consequence of a decrease in the number of connections, but rather reflects a reduction in integration [88] (Fig. 4c), pushing the brain away from the optimal region in connectivity morphospace. Thus, the network dimension may serve as a valuable metric for characterizing pathological shifts in the connectivity morphospace, or the “disconnectivity landscape” [16]. As such, it holds promise as a biomarker for clinical assessment, offering insights into the impairments that result from brain damage.

The local dimension was significantly reduced in regions that became structurally disconnected following stroke (Fig.4f). Rather than simply reflecting spatial proximity to the lesion, this reduction was driven by the extent to which a region lost integrative connections to the rest of the brain. This supports the interpretation that local dimension reflects a region’s capacity to integrate diverse inputs—an ability that is compromised when structural disconnection isolates the region functionally. This interpretation is further supported by our mixed-effects analysis, which showed that graph distance from the lesion—capturing how the impact of a lesion propagates along structural pathways—was a stronger predictor of dimensionality loss than Euclidean (spatial) distance (Fig.4e). Thus, the drop in local dimension is more closely tied to disconnection than to anatomical distance, consistent with the idea that the loss of critical white-matter pathways impairs a region’s capacity for integration [44, 89]. From a functional perspective, this loss in dimensionality may reflect a decline in the flexibility and adaptability of local processing, which could contribute to the cognitive and behavioral deficits observed in stroke patients [90–92].

Indeed, we observed a negative correlation between the dimensionality of stroke-affected connectomes and performance on tasks involving memory and visuospatial skills (Fig. 4d), indicating that lower global dimension is associated with greater cognitive impairment. In contrast, we found no significant relationship between global dimension and sensory-motor deficits. This is consistent with previous work [55], which showed that both local and whole-brain structural (dis)connectivity patterns were more strongly associated with cognitive dysfunction than with sensory-motor impairments. The absence of motor correlations in both studies highlights a selective sensitivity of structural organization—and, in our case, of global connectome dimensionality—to higher-order cognitive function.

This link between structural disconnection and reduced local dimension has broader implications for understanding how network topology shapes brain dynamics. In models of disordered materials, a reduction in dimensionality can suppress critical phenomena, potentially causing the critical transition to vanish [26]. A similar principle may apply in the brain: earlier studies showed that focal lesions reduce markers of criticality in whole-brain dynamics [44]. Here, we extend these findings by showing that such impairments can be explained through a reduction in the dimension of the connectome (Fig. 5e-f). In stroke patients, this lower dimensionality was associated with a marked decline in metastability and an increase in dynamical fragmentation. Metastability has been linked to optimal information processing capabilities, cognitive flexibility, and memory capacity [59, 93, 94], and this interpretation is supported by prior computational [88, 95] and empirical studies [44, 96, 97]. Therefore, the dimensionality of the connectome appears to be a crucial factor in understanding impaired information processing and the related cognitive deficits.

### E. Limitations and future perspectives

The relation between structural dimension and criticality was investigated with the aid of a computational model. While these models have proven to be reliable tools for reproducing functional activity both in health and disease [13, 98], they may not fully capture the dynamical mechanisms occurring in real brains. Future research should focus on directly comparing network dimensionality with empirical functional data to further validate the obtained insights. Additionally, it would be valuable to investigate how these relationships hold across different neurological and psychiatric conditions and during normal aging. In this context, it would be intriguing to investigate the potential disruption of criticality occurring in pathologies characterized by structural overconnectivity (rather than disconnection), such as autism spectrum disorder.

## IV. METHODS

### A. Estimation of network dimension

To estimate the dimension of a network, we apply the algorithm proposed in Peach et al. [39]. Briefly, this model capitalizes on the temporal response of impulsive perturbations that diffuse on graphs to infer a nodal measure of dimensionality. In a *d*-dimensional lattice, the time evolution of a localized perturbation placed at position **x**_0_ is

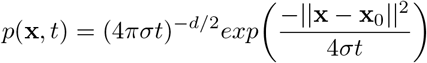

that is the solution of the diffusion equation given *p*(**x**, 0) = *δ*(**x** − **x**_0_) as initial condition and *σ* as a diffusion coefficient. The transient response display a maximum of amplitude 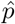 at time 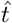 in any other location **x** given by [49]

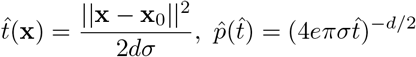

Such values can be then used to infer the dimension of the lattice as

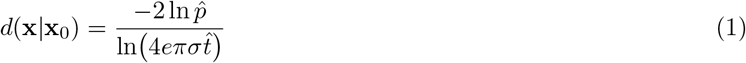

The same recipe can be generalized on graphs. Given a network described by an adjacency matrix *W*, the temporal evolution of the time-dependent node vector **p**(*t*) follows

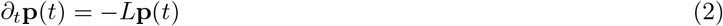

where *L* = *K*^*−*1^(*K* − *W*) is the (normalized) graph Laplacian, and *K* is the diagonal matrix of node degrees. By perturbing the node *i* with intensity *δ*_*i*_, such that **p**(0) = (0, 0, …, *δ*_*i*_, …, 0), the response of node *j* is given by *p*_*j*_(*t* | *i*) = *δ*_*i*_(*e*^*−Lt*^)_*ij*_. Therefore, by numerically simulating (2) and measuring the amplitude 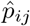 and the time 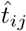 at which a maximum appears, a (relative) dimension can be computed similarly to (1) as

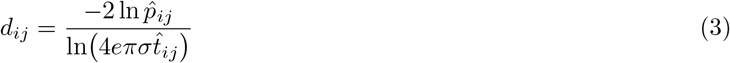

Differently from Euclidean spaces, where the dimension is independent of the starting and observing point (and is always equal to *d*), in complex networks the response is heterogeneous and typically changes between nodes. Moreover, for some pairs of nodes the peak in transient response may be even missing due to boundary effects [49]. Thus, the local dimension of a node *i* at a given scale *τ* can be defined by averaging the relative dimension of nodes displaying a peak in their response before a given time *τ*

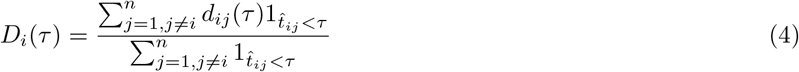

and, finally, the global dimension of the network as 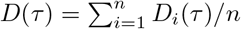.

Following previous work [39], we identify the optimal scale *τ* at the peak of the global dimension *D*(*τ*), as this is the scale where the framework’s results match theoretical predictions in Euclidean lattices. For empirical connectomes, where no theoretical expectation exists, this provides an a priori choice of scale, ensuring a consistent and statistically robust framework for comparison across the cohort. The robustness of this criterion is further supported by the broad and smooth shape of *D*(*τ*) across all subjects, with individual peak locations clustering consistently in the range *τ* ≈ 4 − 7 (see Supplemental Information, Fig. S7).

To verify that the observed reduction in dimension after stroke is robust to different estimation methods of network dimension, we compute also the spectral dimension [30, 35, 36] of the connectomes. The spectral dimension *D*_*s*_ of a network can be estimated from the average return probability *R*(*t*) since it is expected to scale as

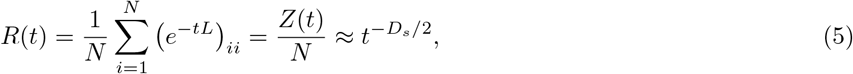

where *Z*(*t*) = Tr *e*^*−tL*^ is the partition function of the associated propagator [42, 63] (see Supplemental Information, Fig. S6 for more details about its computation).

### B. Synthetic networks

To quantify the effect of some paradigmatic topological features on network dimensionality, we generate random networks from different ensembles. Specifically, to investigate the effect of topological disorder, we generate small world networks [45] — that is, models where nodes are initially connected in a lattice-like regular pattern, and then the links are shuffled with a given rewiring probability *p*_*rew*_ to introduce topological shortcuts. Therefore, these networks span from regular topologies with nodes connected only to their nearest neighbors, up to randomized structures.

Instead, to investigate the interplay between segregation and integration on network dimensionality, we consider a stochastic block network [46] — that is, models reproducing groups or communities of nodes that are randomly connected internally with probability *p*_*in*_ and externally with probability *p*_*out*_ — with four communities. This class of models is useful to investigate the mesoscale organization of complex networks. We vary the mixing parameter, defined as 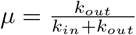, where *k*_*in*_ is the average number of connections a node has with the nodes in its community, and *k*_*out*_ is the average number of external connections. Values of *µ* ≪ 1 generate a highly segregated network with most connections existing only within a community, whereas *µ* ≈ 1 gives a network with no particular community structure, resembling Erdos-Renyi random networks.

Moreover, to account for weight heterogeneity, we generate Erdős-Rényi networks with weights drawn from a lognormal distribution. The parameters of the lognormal distribution, *µ* and *σ*, are selected to vary the distribution’s variance while maintaining a constant average link strength (equal to unity).

Average strength *k* was also systematically varied across all models to disentangle the effect of link strength from the other topological features.

### C. Stroke dataset

We re-analyzed the dataset recently published in [55], which comprises patients with first-time stroke enrolled at Washington University in Saint Louis. Specifically, 79 patients were included (mean age = 60.1 ± 11.5 years; mean education = 13.3 ± 2.5 years). All stroke patients were enrolled in the subacute phase, within two weeks of stroke onset. A healthy control group of *n* = 33 participants was also included, matched to the patients by age (*t* = 0.927, *p* = 0.356), sex (*χ*^2^ = 1.433, *p* = 0.231), and years of education (*t* = −0.897, *p* = 0.372).

Structural connectivity matrices were derived from diffusion-weighted imaging using tractography and parcellated according to the Schaefer atlas at multiple spatial resolutions (100, 200, and 500 parcels). Full details on diffusion preprocessing and tractography procedures can be found in [55]. The dense connectomes were then thresholded to retain the top 20% of connections, to improve the signal-to-noise ratio [99] and to ensure that the two groups had the same network density.

Participants underwent an extensive behavioral battery assessing the following cognitive domains: memory, language, visuospatial attention, executive functions, and motor abilities (see [55] for details). Dimensionality reduction was performed using hierarchical factor analysis, applied separately to stroke patients and healthy controls. This revealed five main latent factors accounting for approximately 50% of the total behavioral variance. The first factor primarily loaded on verbal memory, language, and executive function tests. The second and third factors captured right and left motor function, respectively. The fourth factor reflected visual memory, working memory, and general attention performance, while the fifth factor was mainly associated with lateralized visuospatial abilities. For a full description of the latent behavioral dimensions and their relation to structural disconnection patterns, we refer the reader to [55].

### D. Network integration

We quantify network integration as *I* = 1 − *Q*, where *Q* is the modularity index for a specific network partition. The modularity *Q* is given by

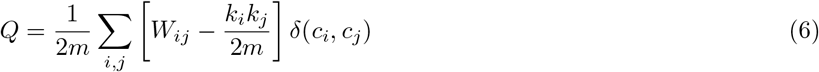

where *m* is the sum of all edge weights in the network, *c*_*i*_ is the community to which vertex *i* is assigned, and *δ*(*c*_*i*_, *c*_*j*_) is the Kronecker delta function, which is 1 if nodes *i* and *j* belong to the same community, and 0 otherwise. Modularity is the fraction of the edges that fall within the given groups minus the expected fraction if edges were distributed at random. Therefore, it measures the extent to which nodes within the same community are more densely connected to each other than to nodes in other communities. High modularity indicates a strong community structure, with more intra-community connections than would be expected by chance. By contrast, a higher integration value implies that communities are more interconnected, with an increased number of links bridging different communities, thus promoting global cohesion within the network.

To define the most representative partition, we used consensus clustering from 1,000 repetitions of the Louvain algorithm [100]. Given that this partitioning method might not yield statistically significant community structure [101], we also identify network partitions via statistical inference using the stochastic block model (SBM) [46, 102] (see Supplemental Information, Fig. S3).

While network integration can be assessed using other metrics (such as network entropy or global efficiency - see [44]), the choice of modularity is motivated by its intuitive interpretation and its direct manipulability in generative models, such as those used for our synthetic networks. To further validate this choice empirically, we assessed the relationship between modularity *Q* and global efficiency across all subjects, finding a significant negative correlation (*ρ* = −0.34, *p* = 0.003; see Supplemental Information, Fig. S13), supporting the use of 1 −*Q* as a proxy for integration in connectome-like networks.

### E. Stochastic whole-brain model

To simulate neural activity at the individual level we employed the whole-brain stochastic model introduced in [56] to describe the dynamics of the human brain at a mesoscopic scale. Such a model is a variation of the Greenberg & Hastings cellular automaton [103]. Briefly, each node in the system belongs to one of three states: quiescent *Q*, excited *E*, or refractory *R*. The original dynamics of the GH automaton is modified in such a way that the states undergo the following stochastic transitions:

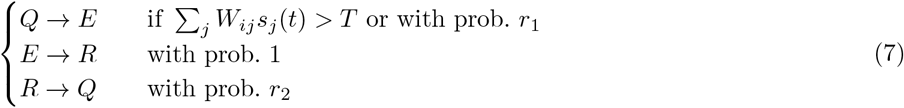

where *s*_*j*_(*t*) ∈ {0, 1} is the state of node *j* at a certain time step *t* - set to 1 if the node is in the *E* state, and 0 otherwise -, *W*_*ij*_ is the weighted connectivity matrix of the underlying network, *r*_1_ is the probability of self-activation and *r*_2_ is the probability of recovery from the refractory state. In particular, *T* is a threshold that governs the induced activation due to interaction with neighboring nodes, which acts as a control parameter of the model. Importantly, we include homeostatic plasticity in the model, implemented as a normalization of the excitatory input of the incoming node 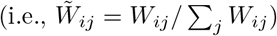. It has been shown that its addition improves the correspondence between simulated neural patterns and experimental brain functional data [57].

Following previous works [44, 57, 58], we set the model parameters to the following values, 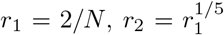, and we vary the activation threshold. We updated the network states *s*_*i*_(*t*) for a total of *t*_*s*_ = 10^4^ time steps, starting from a random initial configuration and discarding an initial transient of additional 5 · 10^3^ time steps. All reported quantities were averaged over 30 independent simulation runs with different random initial conditions.

To characterize the simulated brain activity, for each value of threshold *T* we computed: the mean network activity, 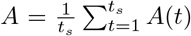, where is the instantaneous activity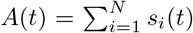; the metastability, i.e., the standard deviation *σ*(*A*) of the average activity *A*(*t*); the average sizes of the largest *S*_1_ and the second largest cluster *S*_2_, called fragmentation. Clusters were defined as ensembles of nodes that are structurally connected to each other and simultaneously active. The value of the critical threshold can be numerically identified by the peak in metastability and fragmentation.

### F. Statistical analysis

To assess the significance of correlations between brain maps while accounting for spatial autocorrelation, we employed the spin test, a non-parametric permutation method [51, 104]. First, each brain map was projected onto a spherical surface, approximating the brain as a sphere for this analysis. We then performed random rotations of the spherical maps, preserving the spatial structure but altering the spatial alignment between the maps. For each rotation (*N*_*spin*_ = 10^4^), we computed the correlation between the rotated map and the other map of interest, resulting in a distribution of correlation values under the null hypothesis of no specific alignment. The observed correlation between the unrotated maps was compared to this null distribution to determine statistical significance.

To investigate the relationship between network integration and the dimension of brain connectomes, while controlling for potential confounding effects of node degree, we performed partial correlation analysis. Specifically, we first removed the effects of node degree from both the network integration measure and the dimension by regressing each on node degree. We then computed the correlations between the residuals of these regressions to determine the direct association between network integration and dimension, independent of degree-related variability.

To assess whether the decrease in local dimension in each region of interest (ROI) in stroke patients is predicted by the (network shortest path and spatial Euclidean) distance of each ROI from the lesion center, we employed a mixed linear model. Specifically, we used the following model: Δ*D*_*i*_ ≈ *L*_*i*_ + (1 + *L*_*i*_ | *ID*), where Δ*D*_*i*_ is the change in local dimension of the *i*-th ROI for the j-th patients with respect to the average among healthy controls, *L*_*i*_ is the distance of the *i*-th ROI from the lesion center, and ID is each participant’s unique ID.

## V. DATA AVAILABILITY

The empirical connectivity matrices underlying this study cannot be publicly shared due to data usage restrictions; detailed information about the dataset and analyses is provided in [55]. The local and global dimensionality estimates computed from these connectomes, obtained using the algorithm of [39], are available at https://github.com/gbarzon/ConnectomeDimension and are sufficient to reproduce the results reported in this manuscript. Group-averaged T1w/T2w map [50] is publicly accessible via the *neuromaps* toolbox [51].

## VI. CODE AVAILABILITY

Custom Python scripts used for all the analysis and simulations are available at https://github.com/gbarzon/ConnectomeDimension. Local and global dimensions were computed using the *DynGDim* package (https://github.com/barahona-research-group/DynGDim). Spin test was performed using the *neuromaps* toolbox (https://github.com/netneurolab/neuromaps). The SBM inference was performed using the *graph-tool* module (https://git.skewed.de/count0/graph-tool).

## ACKNOWLEDGMENTS

We acknowledge R. Leone for a careful reading of the manuscript and valuable suggestions.

