## Supplemental Information for "Structural connectome dimension shapes brain dynamics in health and disease"

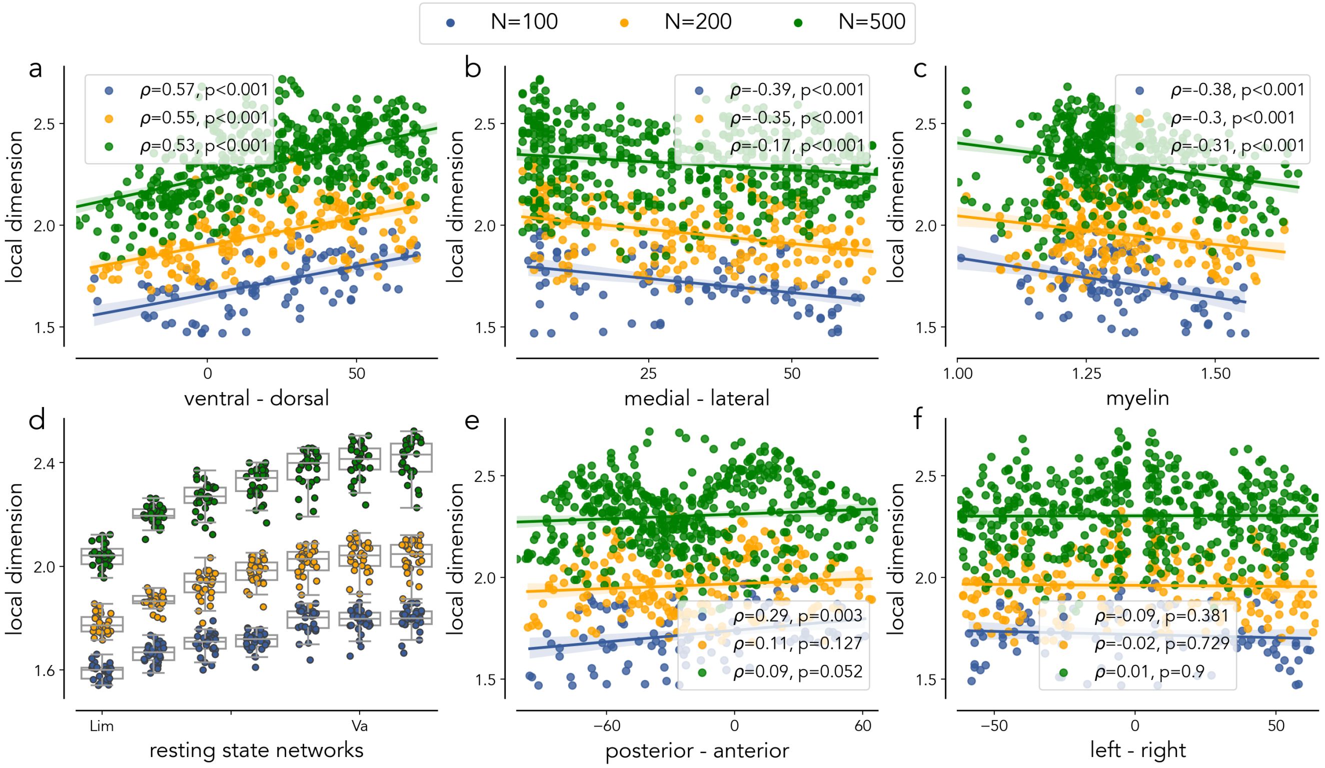


**Figure S1.** Robustness of the relationship between local dimension and brain features across different parcellation schemes. This figure shows the analysis of local dimension for control subjects using coarser (100 nodes) and finer (500 nodes) parcellations, in addition to the 200-node parcellation scheme presented in the main text. (a–d) Replication of the main text analyses: (a) Relationship between local dimension and the ventral-dorsal axis, (b) Relationship with the medial-lateral axis, (c) Correlation with regional myelin content, and (d) Average local dimension within canonical resting-state networks. (e–f) Non-significant relationships between local dimension and the anterior-posterior axis (e) and right-left axis (f).

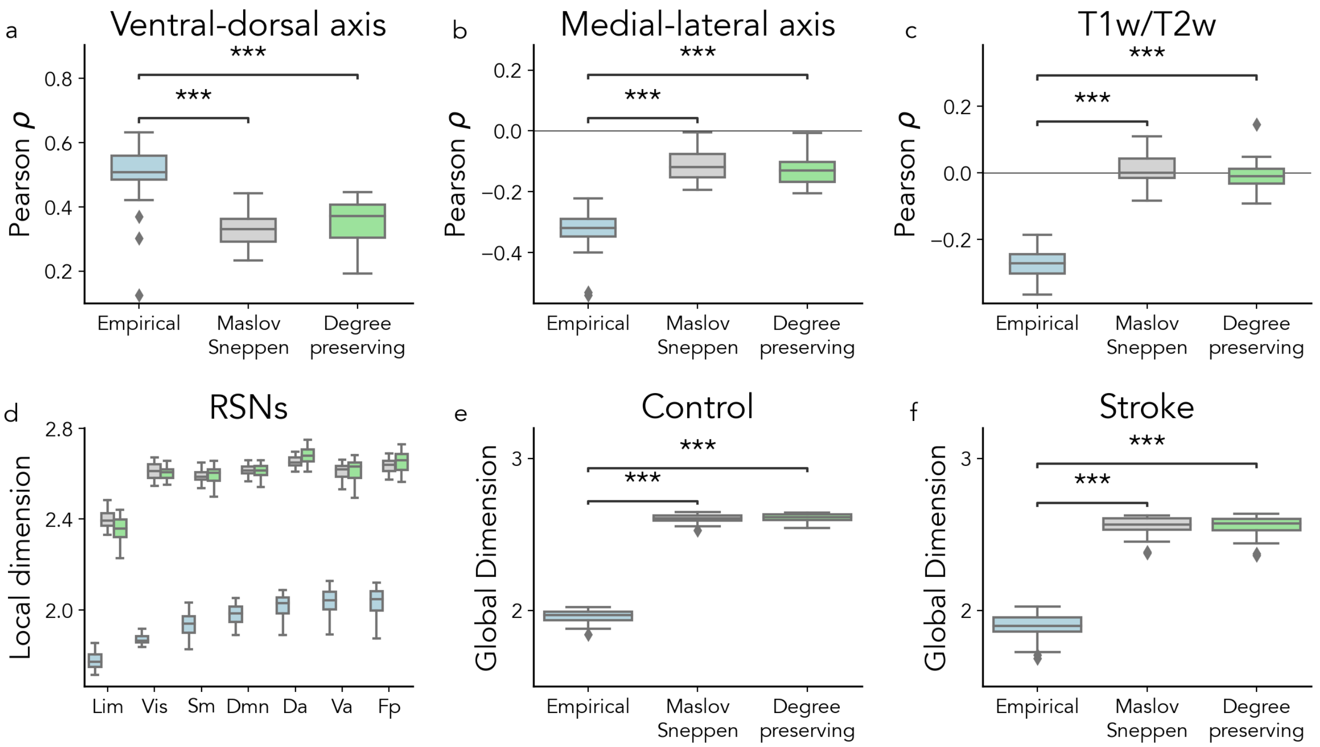


**Figure S2.** Robustness of local and global dimensionality against randomized models. To assess the specificity of dimensionality to the topological organization of the human connectome, empirical data were compared against two null models: Maslov-Sneppen rewiring and degree-preserving randomization. (a–c) Pearson correlation between local dimension and established neurobiological axes, including the ventral-dorsal axis (a), medial-lateral axis (b), and T1w/T2w ratio (c). In all cases, the empirical spatial gradients are significantly altered than those observed in randomized networks ($p < 0.001$, Wilcoxon test). (d) Local dimension across canonical Resting-State Networks (RSNs). The empirical connectome exhibits a clear hierarchical organization (e.g., lower dimensionality in the Limbal network), whereas randomized models collapse toward a higher-dimensional, more uniform state. (e–f) Global dimension for Control (e) and Stroke (f) groups. Empirical global dimensionality is significantly lower than that of null models ($p < 0.001$), demonstrating that the human connectome is organized into a lower-dimensional topological subspace than expected by node degree or weight distribution alone.

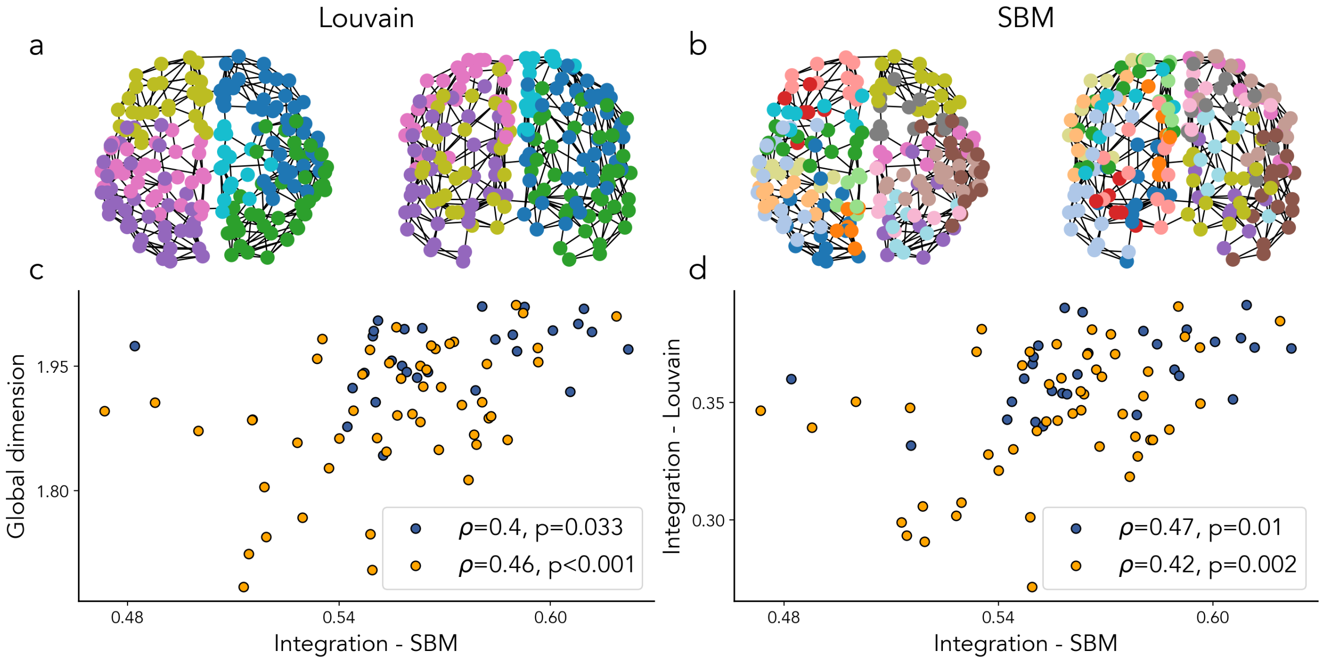


**Figure S3.** Robustness of the relationship between integration and global dimension and their decrease in stroke subjects across community detection methods. (a) Example communities detected in the connectome of an example subject parcellated into 200 nodes using the Louvain algorithm (6 communities detected). (b) Communities detected by inferring a stochastic block model (SBM) from the same connectome (19 communities detected). First, the degree-corrected SBM's description length is minimized by an agglomerative multilevel Markov chain Monte Carlo (MCMC) algorithm. After equilibrating the Markov chain, an estimate of the consensus partition is obtained by sampling 10.000 partitions from the posterior distribution. (c) Significant correlation between global dimension and integration is observed when integration is computed using SBM-based partitions, similar to the results using Louvain partitions shown in the main text. Integration is computed as $I = 1 - Q$, where $Q$ is the modularity of the estimated partitions. (d) Correlation between integration values computed with Louvain-based and SBM-based partitions.

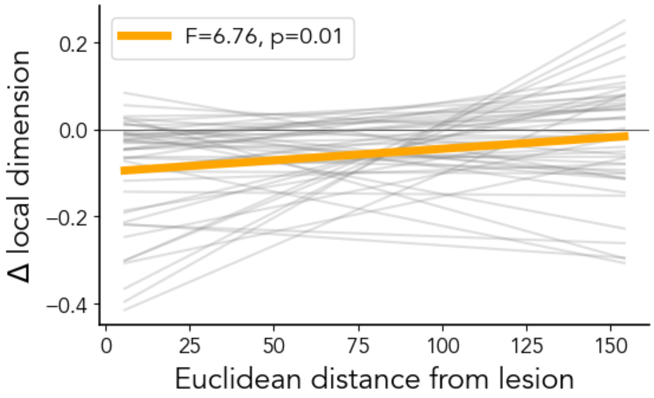


**Figure S4.** Relation between local reduction in dimensionality and Euclidean distance from the lesion, as shown by the random, subject-wise (gray), and fixed, group (orange), effects.

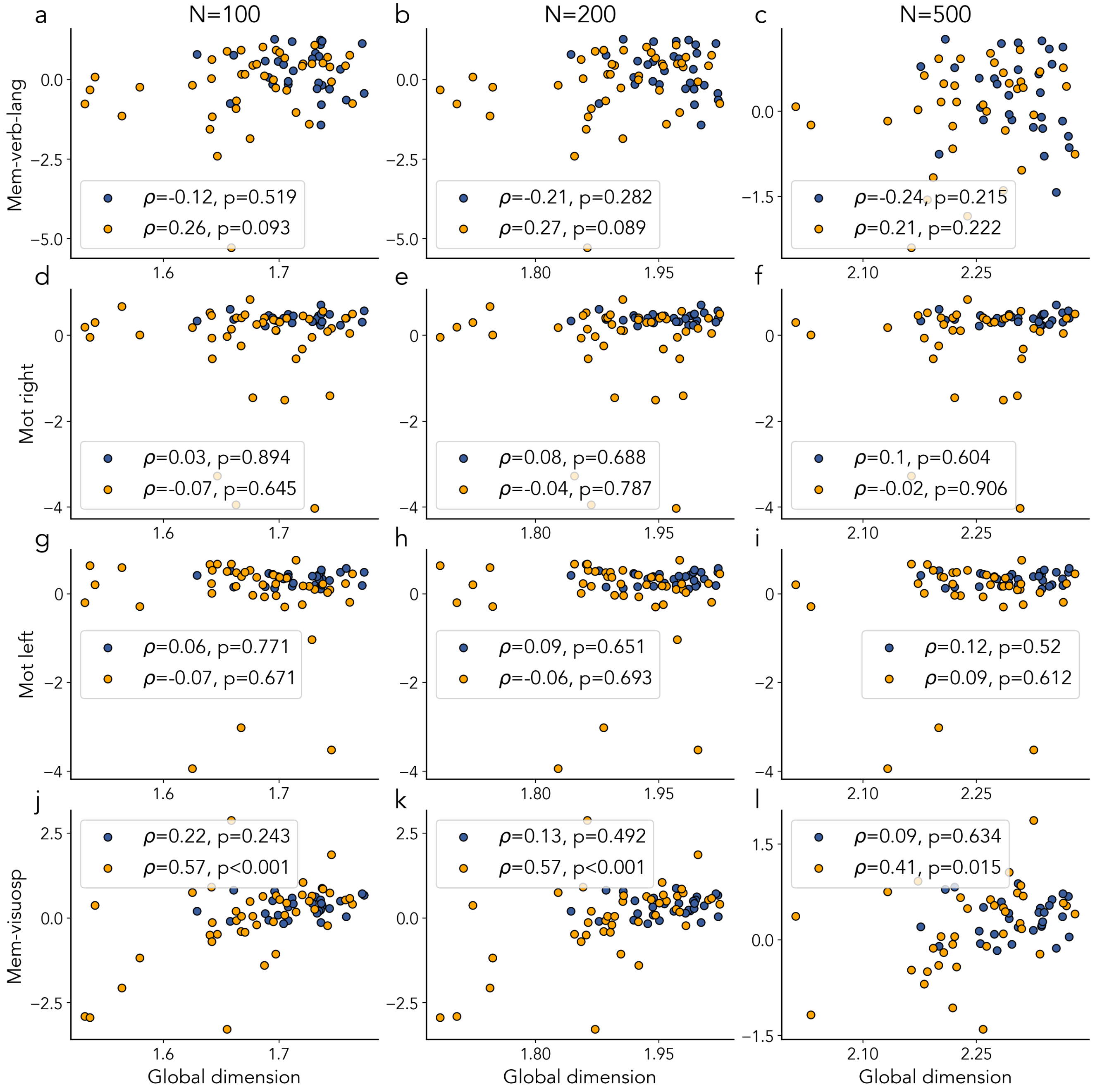


**Figure S5.** Relation between global dimension and behavioral factors across parcellation scales. This figure shows the correlation between global dimension and four behavioral factors—memory-verbal-language (a–c), motor-left (d–f), motor-right (g–i), and memory-visuospatial (j–l)—across three parcellation scales (100, 200, and 500 nodes). Among the factors, only the memory-visuospatial factor shows a significant correlation with global dimension in stroke patients.

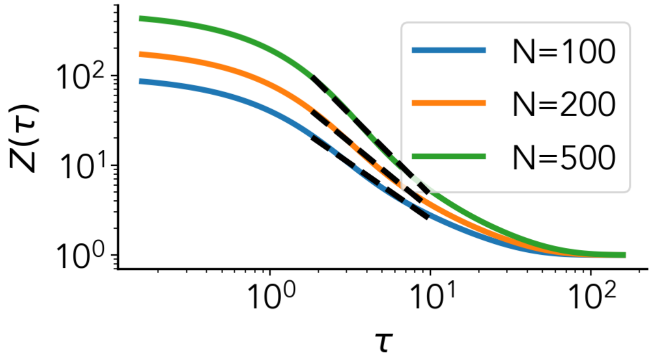


**Figure S6.** Estimation of spectral dimension $D_s$ from the average return probability. The spectral dimension $D_s$ of the network is estimated by analyzing the scaling behavior of the partition function $Z(\tau) = \mathrm{Tr}\left(e^{-\tau L}\right)$, which is computed numerically as a function of $\tau$. This figure shows $Z(\tau)$ for the connectome of one subject at different parcellation scales, plotted in log-log scale. A linear fit is applied to the range in which $Z(\tau)$ exhibits a linear trend on the log-log plot, allowing for the extraction of $D_s$ from the scaling relationship $Z(\tau) \approx \tau^{-D_s/2}$.

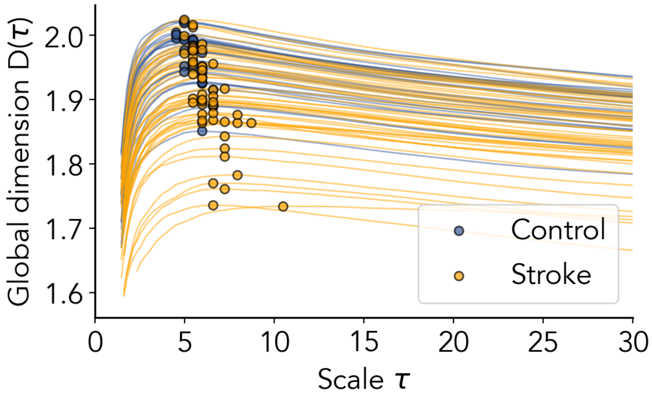


**Figure S7.** Robustness of the scale parameter $\tau$ selection. Global dimension $D(\tau)$ as a function of scale $\tau$ for all subjects. Dots indicate the peak of $D(\tau)$ for each individual, corresponding to the selected scale $\tau$ for the estimation of dimensionality of empirical connectomes. The peaks are broad and smooth across all subjects, indicating that the measure is robust to small perturbations in $\tau$. Individual peak locations cluster consistently in the range $\tau \approx 4-7$, supporting the stability of the criterion across the cohort.

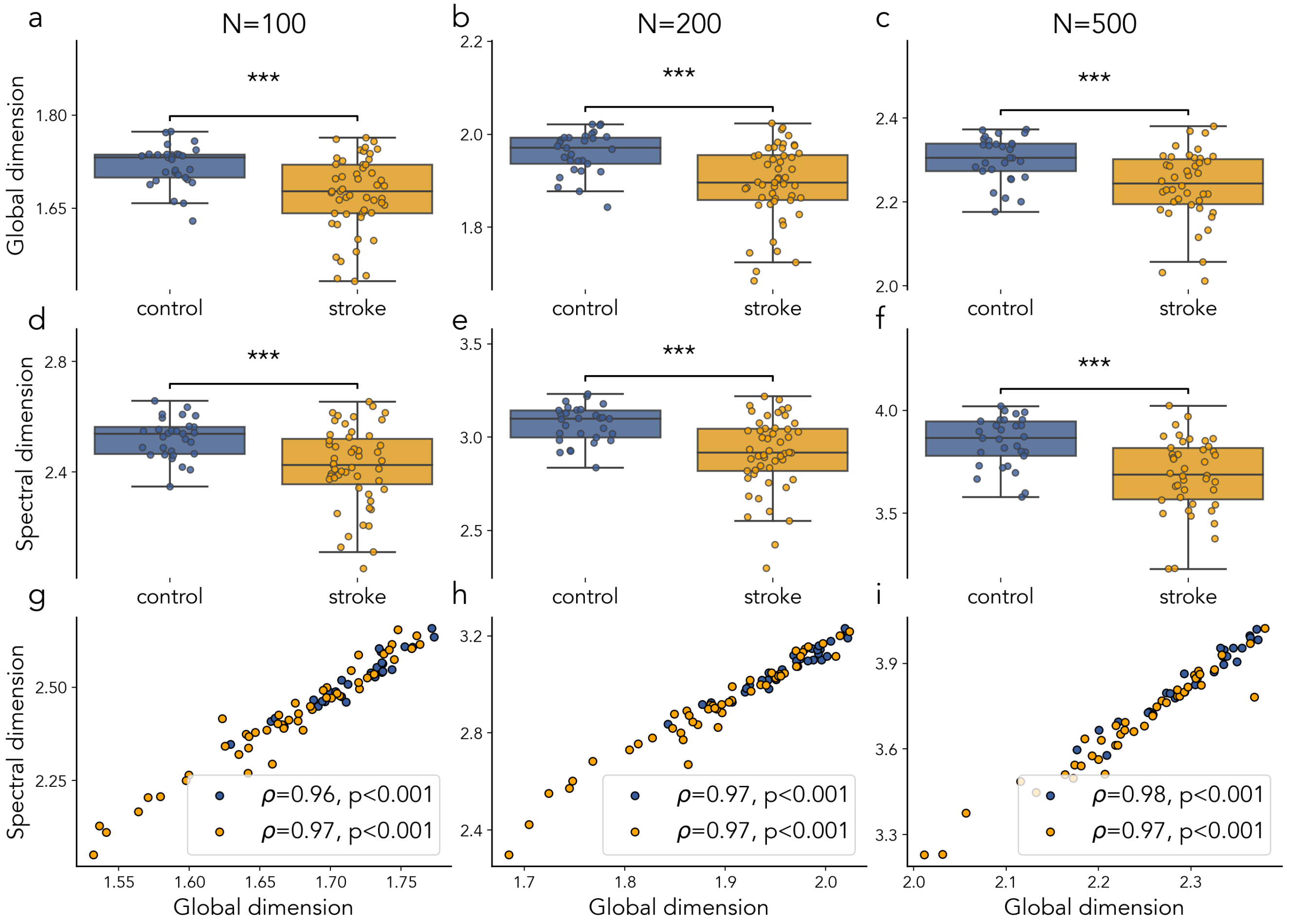


**Figure S8.** Robustness of global dimensionality decrease in stroke across estimation methods and parcellation scales. (a–c) Boxplots of global dimension for control and stroke groups across three parcellation scales (100, 200, and 500 nodes), consistently showing a significant decrease in dimension in stroke patients across both finer and coarser parcellations. (d–f) Boxplots of spectral dimension for control and stroke groups, similarly confirming the reduction in dimension in stroke patients across all scales. (g–i) Correlations between global dimension and spectral dimension across parcellations, illustrating a strong correspondence between the two measures.

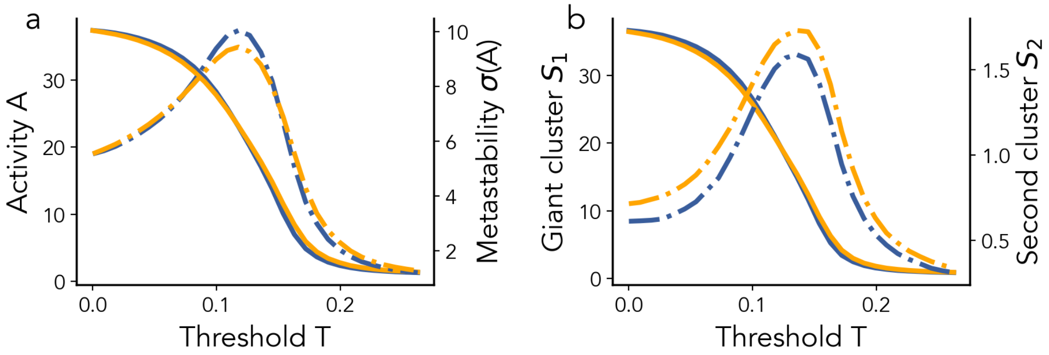


**Figure S9.** Determination of critical observables from simulated brain activity. (a) The average network activity $A$ and its standard deviation (metastability, $\sigma(A)$) as functions of the activation threshold $T$, based on simulations using the whole-brain stochastic model of neural dynamics. (b) Average sizes of the largest cluster $S_1$ and the second-largest cluster (fragmentation) $S_2$ of active nodes, plotted across values of $T$. The critical threshold $T$ is identified where both metastability and the size of the second-largest cluster $S_2$ peak.

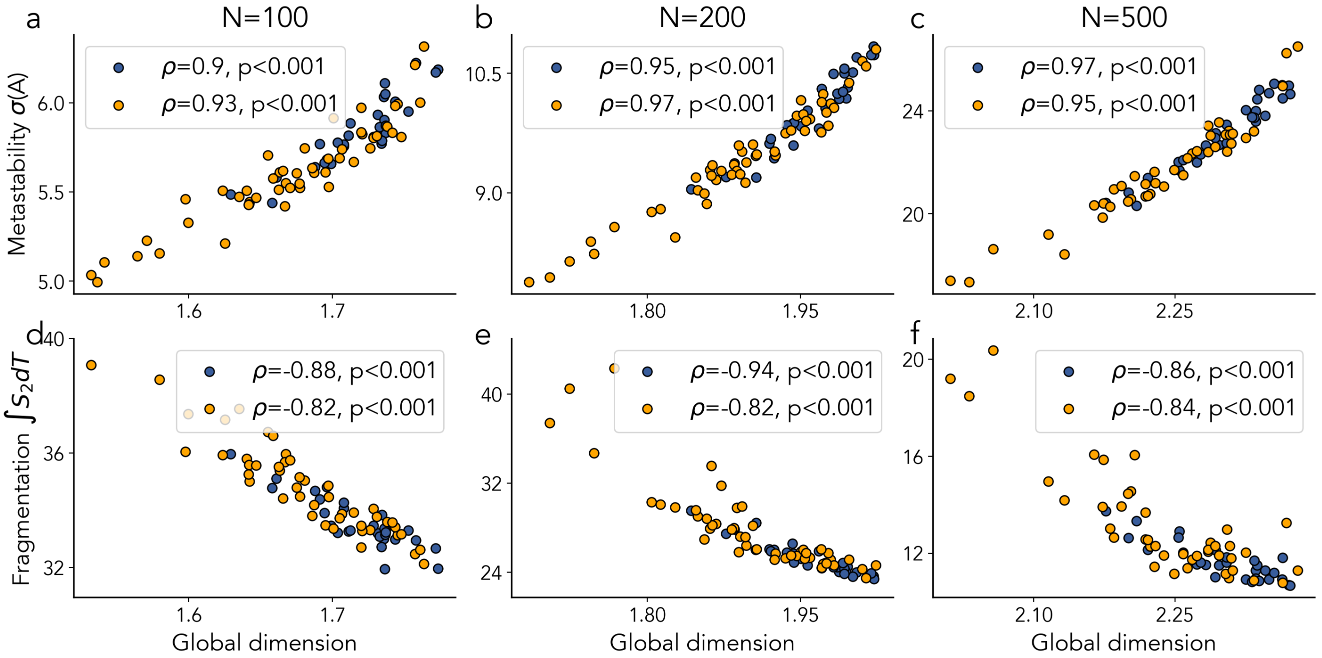


**Figure S10.** Relation between critical dynamics and global dimension across parcellation scales. The first row (panels a–c) shows the correlation between metastability and global dimension for both healthy and stroke groups across different parcellation scales (100, 200, and 500 nodes). The second row (panels d–f) displays the correlation between fragmentation and global dimension for the two groups across the same parcellation scales.

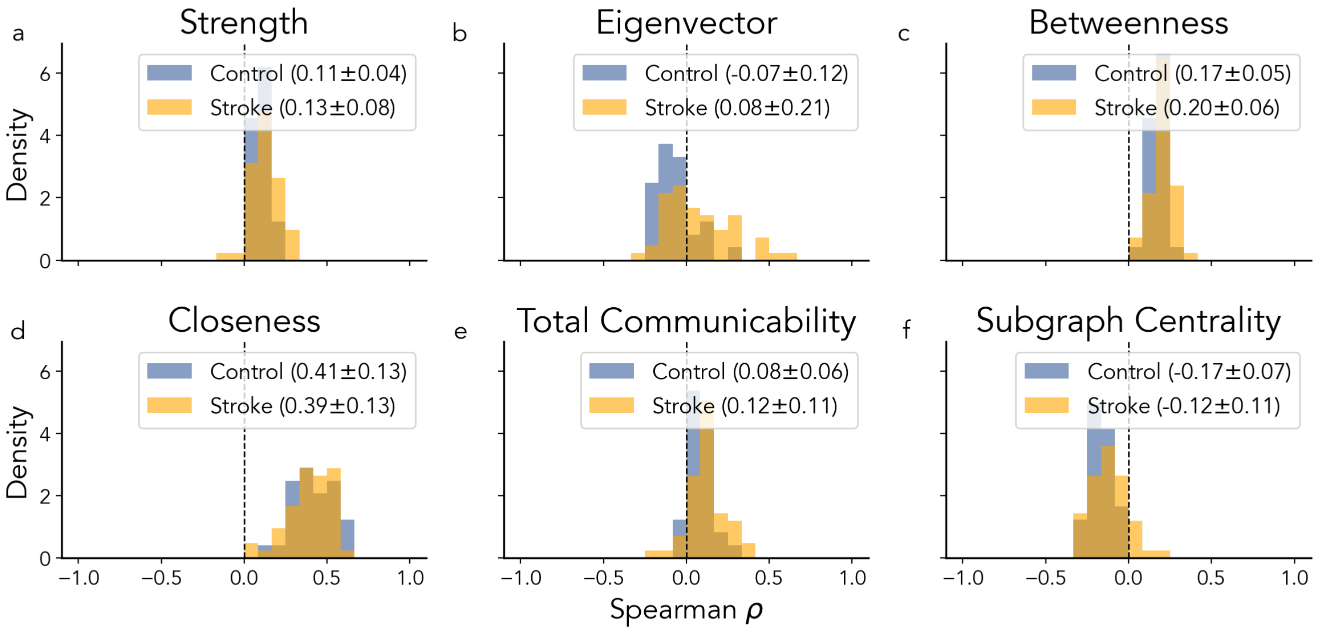


**Figure S11.** Specificity of local dimension. To demonstrate that the local dimension captures higher-order topological features distinct from standard graph-theoretical measures and random configurations, we performed a series of comparative analyses. (a–f) Density distributions of Subject-wise Spearman correlations $\rho$ between local dimensionality and various nodal centrality measures, including Strength (a), Eigenvector Centrality (b), Betweenness (c), Closeness (d), Total Communicability (e), and Subgraph Centrality (f). Correlations are shown for both Control (blue) and Stroke (orange) cohorts. The relatively low and heterogeneous correlation values across these metrics indicate that local dimensionality provides unique structural information not fully captured by established centrality measures.

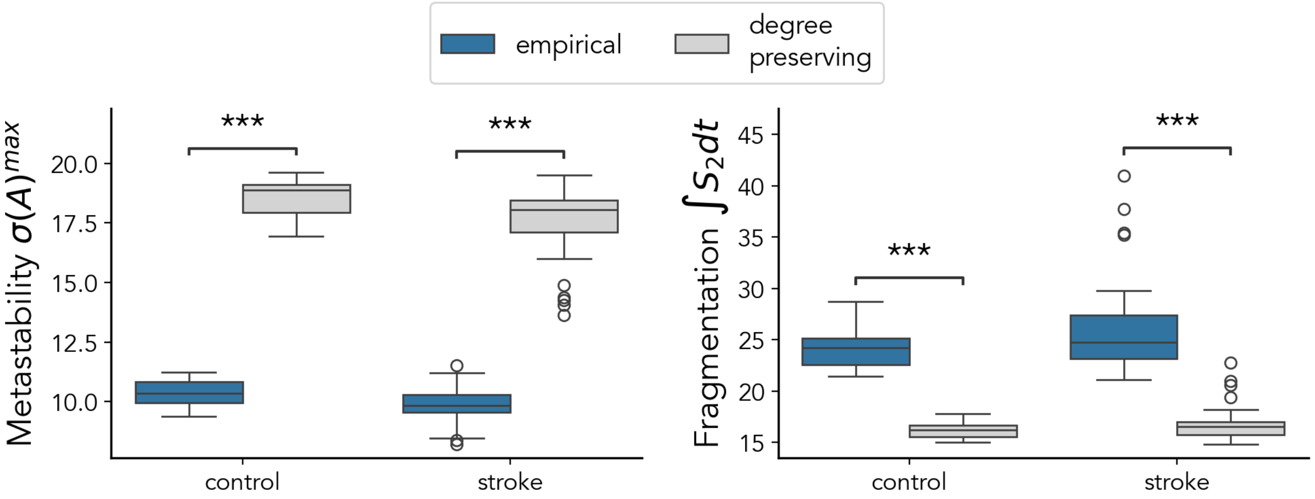


**Figure S12.** Specificity of dynamics under degree-preserving randomization. Comparison of metastability (left) and fragmentation (right) between empirical connectomes (blue) and degree-preserving randomized connectomes (gray), for both control and stroke groups. Degree-preserving randomization significantly alters the dynamical order parameters in both groups, indicating that the observed dynamics are specific to the empirical wiring organization.

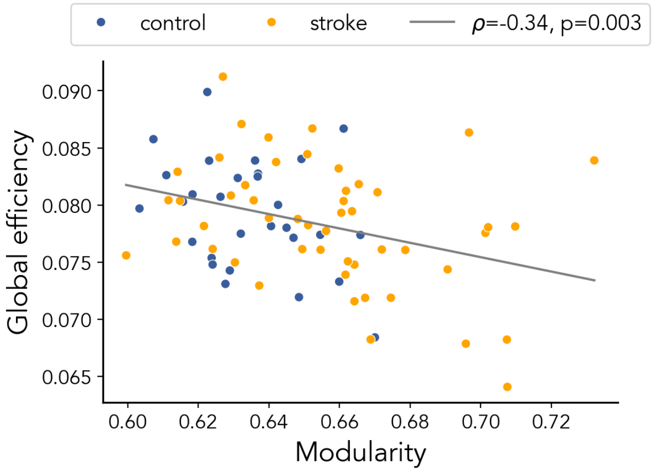


**Figure S13.** Relationship between modularity and global efficiency. Scatter plot of modularity $Q$ versus global efficiency for all subjects (controls in blue, stroke patients in orange). The gray line shows the linear fit across all subjects. A significant negative correlation is observed , indicating that lower modularity (i.e., larger integration) is associated with higher global efficiency.
